# Commentary on “Limitations of GCTA as a solution to the missing heritability problem”

**DOI:** 10.1101/036574

**Authors:** Jian Yang, S. Hong Lee, Naomi R. Wray, Michael E. Goddard, Peter M. Visscher

## Abstract

In a recent publication entitled “Limitations of GCTA as a solution to the missing heritability problem” Krishna Kumar et al. (2015 PNAS) claim that “GCTA applied to current SNP data cannot produce reliable or stable estimates of heritability”. Here we show that those claims are false and that results presented by Krishna Kumar et al. are in fact entirely consistent with and can be predicted from the theory underlying GCTA.

---

GCTA, more precisely, the GREML approach (2) implemented in the GCTA software tool (3), is developed to estimate the total genetic variance in a trait explained by the SNPs that have been genotyped in an experiment. It is not assumed that these SNPs actually cause variation in the trait. Rather it is assumed that the SNPs are in linkage disequilibrium (LD) with the causal variants and therefore tag them to some degree. No assumption is made in GREML that SNPs are in linkage equilibrium. As the number of SNPs used is increased, the extent to which they tag the causal variants through LD may increase so that the total genetic variance explained by the SNPs increases, although the variance explained per SNP may decrease simply because the total variance is spread across more SNPs. In Yang et al. (2), we presented theory and experimental results to show how the total variance explained by the SNPs increases towards a plateau as the number of SNPs used is increased. The theory is based on a statistically equivalent model of fitting effects of SNPs and genome-wide effects of individuals as random effects and indeed addresses 'missing heritability' by estimating the total variance that would be explained from genome-wide significant SNPs in an infinite sample of individuals using the same SNP chip. The estimate from GCTA-GREML is distinct from a pedigree-based estimate of genetic variance because a pedigree-based estimate is independent on how much of total variance is tagged by the SNP chip. Since our original publication in 2010, there have been a series of method developments in estimating genetic variance from SNP data in unrelated individuals for human complex traits or common diseases (4-8).

Krishna Kumar et al. appear to misunderstand the assumptions of GCTA-GREML with a statement that “GCTA assumes that the SNPs used are in linkage equilibrium” (in their page 2). They therefore mistakenly believe that the variance explained per SNP should be the same regardless of the number of SNPs fitted in the model. This is not the case and is not an assumption of the method. In fact, GREML fits all the SNPs jointly in a random-effect model so that each SNP effect is fitted conditioning on the joint effects of all the other SNPs (i.e. it accounts for LD between the SNPs). This is analogous to a linear regression analysis (fixed-effect model) of multiple SNPs. The difference is that in the analysis of a random-effect model we assume the SNPs effects (fitted jointly) following a normal distribution so that the model is solvable even when *m* > *N* (*m*= number of SNPs and *N* = sample size). The variance of SNP effects (denoted as 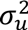 in our original publication (2) and *σ*^2^ in Krishna Kumar et al.) is interpreted as the variance of a SNP effect when it is fitted jointly with all the other SNPs. We show below that *σ^2^* is constant if all the SNPs are in linkage equilibrium (LE), i.e. independent, and that *σ*^2^ for a random subset of SNPs is larger than that for the entire set if SNPs are in LD.

Krishna Kumar et al. argue that GCTA-GREML gives incorrect estimates because it overfits the data since the number of SNPs used is usually greater than the number of individuals with phenotypes. This would be true if SNP effects were estimated as fixed effects. As we have mentioned above, GREML fits all SNPs jointly in a random-effect model. Such models are widely used, for instance in livestock genetics (9, 10), where it is common place to fit models in which the number of random effects exceeds the number of animals with records. The only parameters estimated in a REML analysis are the variance components, the number of which is typically much smaller than sample size, so there is no over-fitting.

Krishna Kumar et al. observed from simulations of unlinked SNPs (shown in their Figure 2) that the sampling variation of 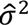 for a random subset of SNPs was much larger than that for the entire set. Their claim that therefore this is a failure of GCTA-GREML is incorrect because the sampling variance of 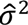 is expected to increases with decrease in number of SNPs. Following Visscher et al. (11), the sampling variance of the estimate of *V*_g_ is var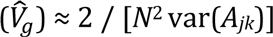, where *A_jk_* denotes the off-diagonal elements of the genomic relationship matrix (GRM). If SNPs are in linkage equilibrium (LE), var(*A_jk_*) = 1 / *m* (2, 12). We then have var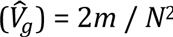. Therefore, 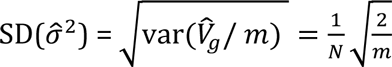. For *N* = 2,000, this equation predicts that 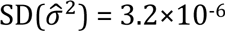 when *m* = 50,000, highly consistent with the observation in Krishna Kumar et al. of 3.1χ10^−6^. This equation also predicts that 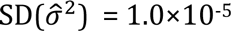 when *m* = 5,000. Therefore, the 95% confidence interval (CI) of the distribution shown in their Figure 2 should be 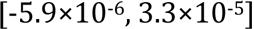, rather than 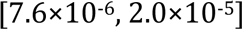 as indicated by the red arrows. Note that Krishna Kumar et al. used the default setting of GCTA so that all the estimates were constrained to be positive. There is an option in GCTA (—reml-no-constrain) that allows negative estimates so that the mean from multiple replicates is unbiased. Our predicted 95% CI again fits well with their observed results in Figure 2. Hence the results from their analyses are expected and consistent with the theory underlying GREML.

Krishna Kumar et al. simulated 50,000 unlinked SNPs so that *σ*^2^ for a random subset of 5,000 SNPs 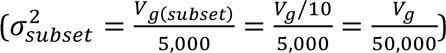 is expected to be the same as that for the entire SNP set 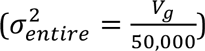. They then claim that this is also the case in real data where SNPs are not independent. This misconception is the essential problem in Krishna Kumar et al., and has led to inappropriate design of their analytical experiments, misinterpretation of the GREML results, and incorrect and confusing inference about the bias in GREML estimates. This is surprising given that the impact of LD on GREML estimation has been explored in depth in recent publications (12-14).

In their Figure 4 (results presented in their Figure 7 are similar), they performed a GCTA-GREML analysis for a real data set (2,698 individuals genotyped on 49,214 SNPs from the Framingham Heart Study, FHS) for systolic blood pressure (SBP). They claimed that the GREML estimates are biased because they observed that 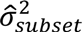 for 5,000 randomly sampled SNPs were on average much larger than 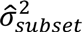 and were widely distributed beyond their calculated 95% CI (indicated by the red arrows in their Figure 4A). They attributed the “bias” to population stratification, and found that fitting eigenvectors from principal component analysis as fixed effects could not fix the “bias” (their pages 3 and 4). As we have mentioned above, 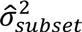 is expected to be larger than 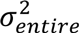 because in the analysis of a subset of SNPs each SNP is fitted conditioning on a much smaller number of SNPs than that in the analysis of the entire set. This is analogous to that in a linear regression analysis of multiple correlated SNPs where the mean variance explained for a subset of SNPs is expected to be larger than that for all SNPs. In an extreme scenario where only one SNP is fitted at a time, the mean variance explained by SNPs estimated from single-SNP analyses is clearly much larger than that from a joint analysis. This also explains why *V_g_* does not decrease linearly with the decreased number of SNPs as observed in Yang et al. (2) and mentioned in Krishna Kumar et al. (they called it “saturation of heritability estimate”). Moreover, if there are related individuals in the sample, 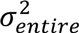 would be even larger because not many SNPs are needed to capture the pedigree relatedness. Krishna Kumar et al. claimed that they used 2,698 unrelated individuals from the FHS data (without describing how these unrelated individuals were selected so that the exact analyses cannot be duplicated by us and other readers), which is much larger than the number of unrelated individuals reported in a previous study (see Supplementary Note 2 of Wray et al. (15)), or the number shown in our Figure 1 using a relatedness threshold of either 0.1 or 0.05 (Note that a threshold of 0.05 is very stringent in this case because of the large sampling error in GRM due to the small number of SNPs used). This suggests that it is very likely that there is remaining (cryptic) relatedness in the FHS data used in Krishna Kumar et al., which would lead to inflation in 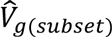 and 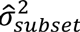. In conclusion, Krishna Kumar et al. used the incorrect expected mean and SD of 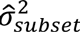 to compare with those observed from resampling, and therefore their conclusion about biasedness of GREML estimates is not supported by empirical evidence. They attributed the “bias” to population stratification, which is also a false statement.

**Figure 1.**
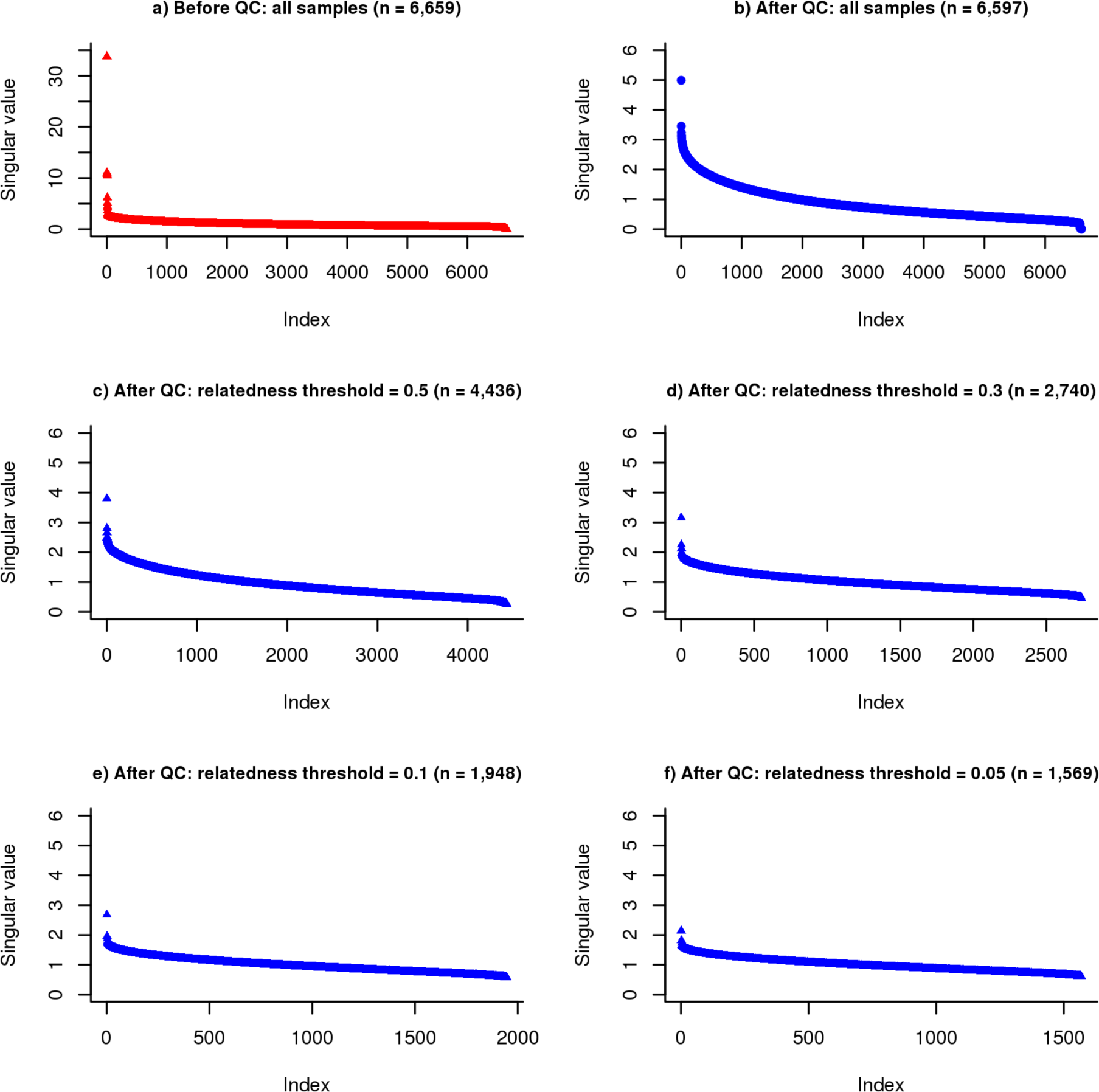
Singular values of SNP genotype matrix estimated from the FHS data. We accessed the FHS data through dbGaP (phs000007.v16.p6). There are 6,659 individuals and 49,094 SNPs before QC. We removed individuals or SNPs with missingness rate > 0.05, and excluded SNPs with Hardy-Weinberg Equilibrium p-value < 0.001 or minor allele frequency < 0.01. We retained 6,597 individuals and 35,221 SNPs after QC. Singular values of the SNP genotype matrix **Z** are the square roots of eigenvalues from a principal component analysis of the GRM. In panels (c), (d), (e) and (f), we used a range of thresholds to remove cryptic relatedness using —grm-cutoff option in GCTA.

Krishna Kumar et al. also claim that GCTA-GREML has a fundamental problem because the singular values of the matrix of genotypes (**Z**) contain values that are close together or are close to zero. However, we could not reproduce these extremely large and small singular values using FHS data also downloaded from dbGaP with or without quality control (QC) on SNP genotypes, or whether filtering the GRM for relatedness (see our Figure 1). Even if there are singular values that are close together or are close to zero, this does not lead to unreliable estimates of the variance explained by the SNPs. Recent theoretical studies (16, 17) have clearly shown that the standard error of the estimated variance is largely dependent on two quantities – the sample size and the variance of the eigenvalues of the sample **ZZ**’ (which is proportional to the GRM), and neither eigenvalues close together or near zero cause large standard errors. The authors then proposed a “denoising” approach by setting the small eigenvalues to zero, which clearly loses information and will lead to underestimation of the variance explained by SNPs. For instance, it is shown in their Figure 6 that the heritability estimate is unbiased without adjustment (Figure 6A; an estimate of 0.62 is not significantly different from the true parameter 0.65 given SE = 0.22) and biased when the GRM is adjusted using the denoising approach (Figure 6B; an estimate of 0.17 is significantly different from 0.65 given SE = 0.22).

In addition to the many incorrect claims about properties and effects of the SNP-derived GRM, Krishna Kumar et al. claim that GREML is sensitive to measurement errors in the phenotypes (their page 4) because they observed variation in GREML estimates when the phenotype of each individual was randomly sampled from repeated measures (their Figure 5). However, such variation is entirely expected if the correlation between repeated measures are not perfect, and is not be specific to a particular method. For example, it also applies to the estimate of a sample mean in a simple predictable way.

Finally, we want to emphasize that GCTA-GREML (2, 3) is a method that was originally proposed to estimate the proportion of variance explained by all SNPs (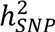) on a SNP genotyping array rather than total (narrow sense) heritability (*h*^2^). The method fits all the SNPs simultaneously. The analysis was strictly limited to unrelated individuals to address the problem of ‘missing heritability’ and to avoid possible confounding from shared environment effects between relatives (2). Further development of the method has allowed for estimating 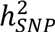 and *h*^2^ simultaneously in family data (18). For completeness, we note that in extension of the GREML method to disease traits more caution is needed compared to analysis of quantitative traits, because any genotyping factors confounded with case-control status could be partitioned into the GREML estimate (4). The GREML estimate of 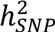 is the lower limit of heritability because it is very unlikely that all the causal variants (in particular those in low minor allele frequency) are all perfectly captured by the SNPs used in GWAS. There has been substantial analytical work demonstrating that 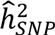 is an unbiased estimate of *h*^2^ if causal variants are a random subset of all SNPs used in the analysis (2, 12-14) and that the reported SE of 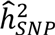 is consistent with the empirical SD of the estimates from resampling (11). Further discussion about how the heterogeneity in LD would impact the biasedness (13, 14) have led to new developments of the GREML method (12), which could be applied to data from whole-genome-sequencing or imputation.

## Acknowledgements

We thank Bill Hill, Alkes Price, John Witte, and Mark Blows for comments and support. This study makes use of the FHS data from dbGaP. The FHS is conducted and supported by the National Heart, Lung, and Blood Institute (NHLBI) in collaboration with Boston University (Contract No. N01-HC-25195). Funding for SHARe Affymetrix genotyping was provided by NHLBI Contract N02-HL-64278. This manuscript was not prepared in collaboration with investigators of the FHS and does not necessarily reflect the opinions or views of the FHS.

